# Stain Normalisation Does Not Consistently Improve Matched-Location Retrieval with Pathology Foundation Models

**DOI:** 10.64898/2026.10.13.758918

**Authors:** Yaning Meng, Binghao Chai, Tapabrata Chakraborti

## Abstract

Pathology foundation models provide reusable representations for computational pathology, but staining- and scanner-induced variation may compromise the robustness and reproducibility of embedding-based analyses across acquisition settings. Stain normalisation is widely used to mitigate such colour variation, yet whether it improves retrieval with modern self-supervised pathology encoders remains unclear. We test whether stain normalisation improves cross-scanner and cross-staining image retrieval using the PLISM whole-slide image subset, comprising 13 co-registered images of serial sections prepared under different staining conditions and digitised using seven scanners (91 whole-slide images). We sample 400 matched locations across the 91 images, yielding 36,400 tile instances. We compare two DINOv2-based encoders, Phikon-v2 (1,024-dimensional) and Midnight-12k (3,072-dimensional), with Reinhard and Macenko normalisation against an unnormalised baseline. Performance is measured using paired top-1 nearest-neighbour retrieval accuracy in two settings: cross-scanner retrieval of the same physical section digitised by different scanners, and cross-staining retrieval across co-registered images of serial sections, which differ in staining condition and may also contain section-level morphological variation. Neither normalisation method consistently improves top-1 retrieval relative to no normalisation, while Macenko reduces accuracy by up to 18.3 percentage points. An exploratory UMAP of unnormalised Midnight-12k slide-level mean embeddings showed partial overlap between scanner groups in a two-dimensional projection. These results challenge the routine use of stain normalisation in foundation-model retrieval pipelines and support validating preprocessing choices for each encoder and retrieval setting.

## 1 Introduction

Foundation models are increasingly used as general-purpose pretrained feature encoders that transform histopathology images into reusable representations for downstream tasks [16, 9]. Many recent encoders use vision-transformer architectures and large-scale self-supervised pretraining, including objectives derived from DINOv2 [11, 16]. Their practical value, however, depends not only on downstream performance but also on model robustness: matched or biologically comparable tissue should remain similar in embedding space despite non-biological variation introduced during slide preparation and digitisation [1, 6, 13].

The appearance of H&E-stained tissue varies across laboratories and imaging systems because of differences in staining protocols, reagent batches, tissue processing, and scanner colour response [14]. Such acquisition-related differences introduce variation that is unrelated to the biological target and can compromise the reproducibility and generalisability of computational pathology models [14, 1, 6]. Stain normalisation methods, such as Reinhard [12], Macenko [7], and Vahadane [15], seek to reduce this variation by transforming source images towards the colour characteristics of a reference image. However, previous evaluations indicate that the effect of normalisation depends on the task, dataset, model, and robustness criterion, and an improvement in one setting cannot be assumed to transfer to another [14, 6]. Whether this preprocessing remains beneficial for pathology foundation models is less clear: pretraining on large, heterogeneous histopathology collections may confer partial tolerance to acquisition-related colour variation, whereas a fixed normalisation transform may interact with learned representations in an encoder-dependent manner [2, 4, 5, 1, 6].

Recent studies have begun to characterise acquisition-related robustness in pathology foundation models. Controlled staining and scanning experiments have demonstrated model-dependent effects on downstream classification tasks [1], while PathoROB showed that embeddings from all 20 evaluated foundation models retained non-biological signatures and that post-hoc robustification reduced, but did not eliminate, associated downstream errors [6]. A co-registered retrieval study likewise reported model-dependent reliability when the same slide was digitised using different scanners [13]. However, these studies do not isolate the effect of classical stain normalisation on paired, matched-location retrieval across both controlled scanner and staining variation, nor establish whether its effect is consistent across encoders [1, 6, 13]. This distinction is important because matched-location retrieval directly tests whether a query and its registered counterpart remain close in embedding space after an acquisition change, without fitting a task-specific classifier [13].

We therefore ask whether colour normalisation improves top-1 retrieval accuracy relative to no normalisation, and whether its effect depends on the encoder or source of appearance variation. We evaluated this question using the registered PLISM subset, which comprises 13 H&E staining conditions digitised using seven whole-slide scanners, yielding 91 whole-slide images [10]. Frozen Phikon-v2 [4] and Midnight-12k [5] encoders are evaluated under unnormalised, Reinhard-normalised and Macenko-normalised settings. Our findings show that, within this matched-location retrieval setting, neither Reinhard nor Macenko normalisation consistently improves retrieval relative to unnormalised inputs, and that the magnitude of the effect varies with the encoder and source of acquisition variation.

## 2 Methods

### 2.1 Dataset

PLISM [10] is a publicly available multi-device and multi-staining histology dataset designed to support controlled evaluation of acquisition-related domain variation. We used the whole-slide-image subset (PLISM-wsi). The dataset contains 13 serial TMA sections, each comprising 46 human tissue types derived from formalin-fixed, paraffin-embedded specimens and prepared under one of 13 different H&E staining conditions. Each section was digitised using seven whole-slide scanners, yielding 13×7 = 91 whole-slide images (WSIs). The digitised images were subsequently co-registered across scanner and staining conditions. We used the preprocessed PLISM tile release provided by Owkin [3], which contains 16,278 matched 224 *×* 224 pixel (0.5 *µm* per pixel) tile coordinates per WSI, stored in a common order across all 91 WSIs. For this study, four contiguous blocks of 100 tile identifiers were selected from this common coordinate list. The same 400 identifiers were retained for every WSI, yielding 400 matched coordinate identifiers and 36,400 tile instances in total.

Two comparison settings were defined for this study. In the cross-scanner setting, the physical section and staining condition were held fixed while the scanner varied, primarily capturing scanner-associated acquisition variation. In the cross-staining setting, the scanner was held fixed while co-registered serial sections prepared under different staining conditions were compared. Because each staining condition corresponds to a physically distinct section, this comparison captures both staining-related differences and unavoidable section-to-section morphological variation and therefore does not isolate staining-related colour variation alone. The dataset subset, comparison design, and matched-location retrieval workflow are summarised in Fig. 1.

**Fig. 1.**
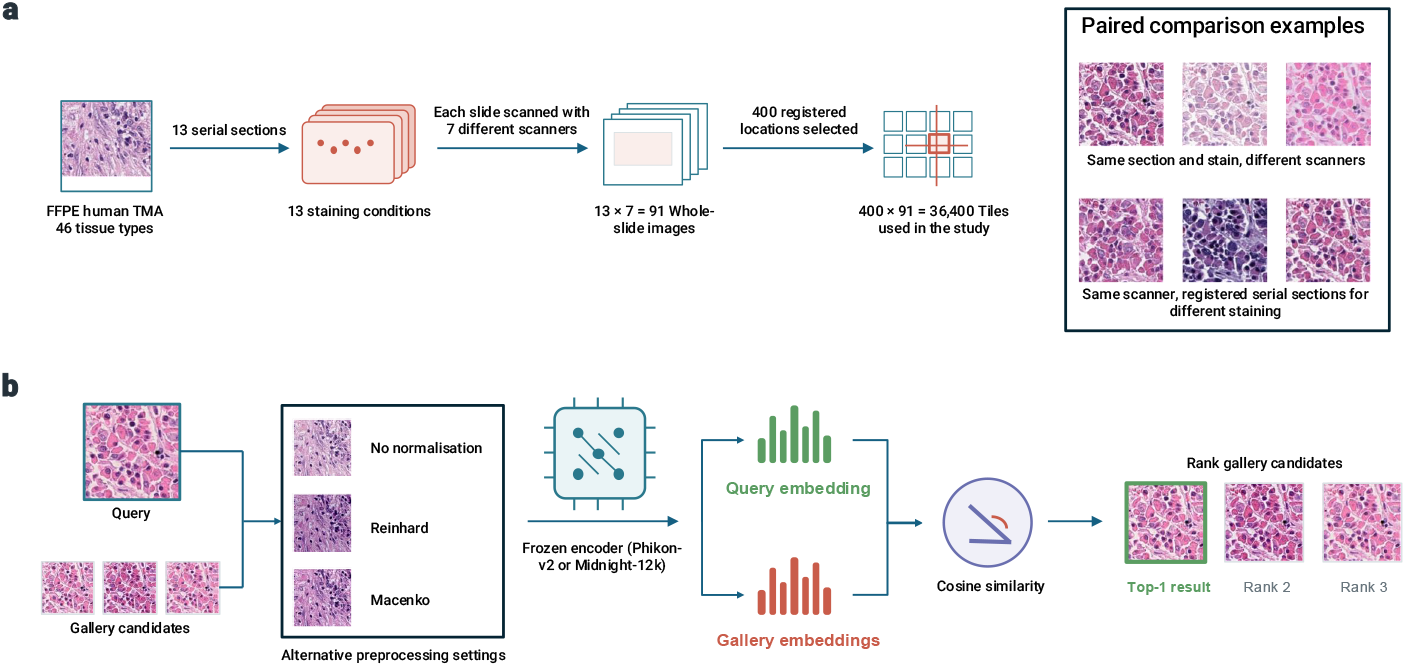
PLISM subset and matched-location retrieval workflow. **a**, Thirteen serial sections from a formalin-fixed, paraffin-embedded human tissue microarray containing 46 tissue types were prepared under different H&E staining conditions; each section was digitised using seven slide scanners, and the resulting images were co-registered. The analysed subset contains the same 400 coordinate identifiers in every WSI, giving 36,400 tile instances across 91 WSIs. Cross-scanner comparisons hold the physical section and staining condition fixed while varying the scanner; cross-staining comparisons hold the scanner fixed while comparing co-registered images of serial sections under different staining conditions. **b**, The same preprocessing setting (no normalisation, Reinhard, or Macenko) is applied to both query and gallery tiles before feature extraction using the selected frozen encoder (Phikon-v2 or Midnight-12k). Gallery candidates are ranked by cosine similarity, and retrieval is correct when the top-ranked tile share the query’s registered coordinate identifier.

### 2.2 Stain Normalisation and Feature Extraction

Tiles were evaluated under three preprocessing settings: no colour normalisation, Reinhard [12] normalisation and Macenko [7] normalisation. Reinhard matches the mean and standard deviation of each channel in Lab colour space to those of a reference tile, whereas Macenko estimates stain vectors in optical density space. Both methods were implemented using torch-staintools [17] and applied identically to query and gallery tiles. A single reference, tile 0 from the GIVH-stained section scanned with the AT2 scanner, was used for all slides. Tiles for which normalisation fails are kept in their original form to preserve paired tile counts; this happens typically near-blank background regions with no recoverable stain signal. Tiles were then encoded using frozen Phikon-v2 [4] (ViT-L/16 architecture; 1,024-dimensional embeddings) or Midnight-12k [5] (ViT-g/14 architecture; 3,072-dimensional embeddings). The two encoders differ in scale and feature construction, enabling us to assess whether normalisation effects are consistent across distinct learned representations.

### 2.2 Evaluation

Retrieval performance is measured by paired top-1 nearest-neighbour accuracy. For a given tile in one condition, we compute its cosine similarity to all tiles in a second condition and record a hit when the top-ranked match corresponds to the same registered location. In the cross-scanner setting this is evaluated over all 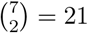 scanner pairs within each of the 13 staining conditions, giving 273 paired comparisons. In the cross-staining setting we evaluate all 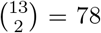 staining pairs within each of the 7 scanners, giving 546 paired comparisons. Results are averaged within each setting.

For exploratory visualisation, the 400 unnormalised Midnight-12k tile embeddings from each WSI were mean-pooled to obtain one slide-level embedding. The resulting 91 slide-level embeddings were projected jointly into two dimensions using UMAP [8], and the same projection was coloured by scanner and staining condition. UMAP was used for qualitative visualisation only and was not treated as a quantitative measure of scanner-associated information.

## 3 Results

### 3.1 Normalisation Does Not Consistently Improve Retrieval

We first asked whether stain normalisation improves tile-level retrieval across scanner and staining variation. Paired top-1 nearest-neighbour accuracy was compared across three preprocessing conditions for both encoders. We evaluated two settings: cross-scanner retrieval compares the same physical section and staining condition digitised by different scanners, whereas cross-staining retrieval compares co-registered images of serial sections prepared under different staining conditions while holding the scanner fixed. In the cross-scanner setting, the unnormalised baseline achieved the highest accuracy for Midnight-12k (56.96%) and was within 3 percentage points of the best Phikon-v2 result. Reinhard normalisation increased Phikon-v2 accuracy from 70.12% to 72.90% but slightly reduced Midnight-12k accuracy, whereas Macenko normalisation produced the largest decline, reducing Phikon-v2 accuracy by 18.30 percentage points to 51.82% and Midnight-12k accuracy by 5.61 percentage points (Fig. 2, panel **a**). In the cross-staining setting, neither normalisation method improved retrieval (Fig. 2, panel **b**): the unnormalised baseline achieved the highest accuracy for both Phikon-v2 (33.78%) and Midnight-12k (29.02%). Accuracy was generally lower in this setting, consistent with the additional morphological differences between co-registered images of serial sections. Overall, the unnormalised baseline matched or exceeded both normalisation methods in three of the four model–setting combinations, indicating that stain normalisation did not provide a consistent benefit for retrieval.

**Fig. 2.**
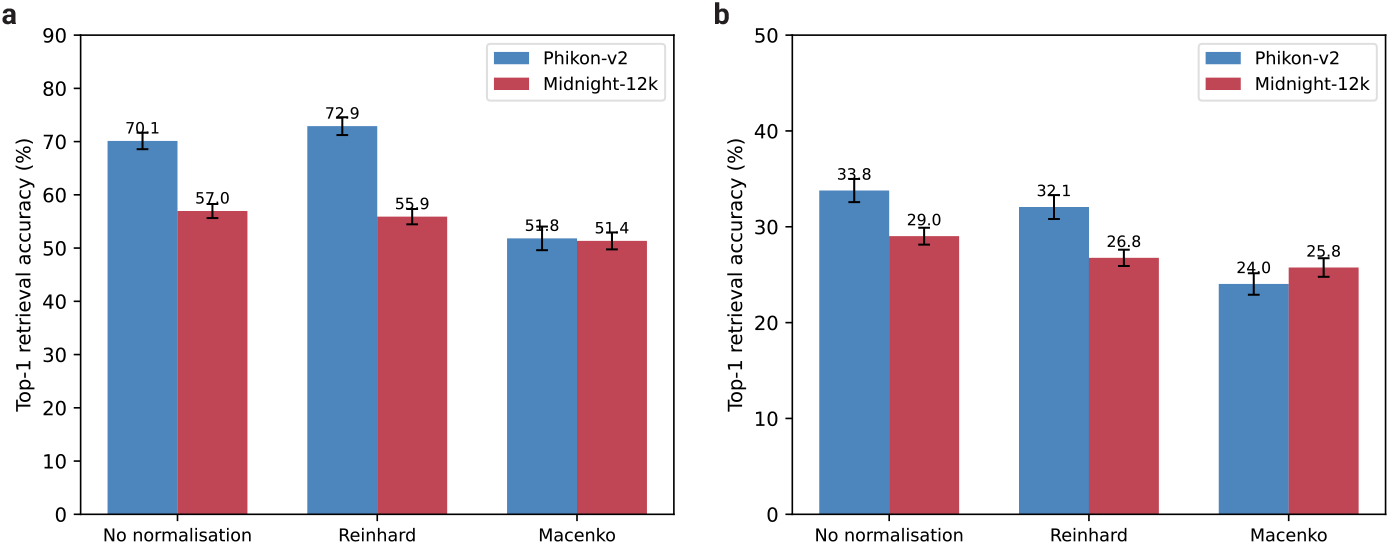
Matched-location top-1 retrieval accuracy for Phikon-v2 and Midnight-12k under three preprocessing conditions. For each unordered slide pair, accuracy was calculated in both retrieval directions, each using 400 query tiles and 400 gallery candidates, and the two directional accuracies were averaged to obtain one pair-level value. Bars show the mean pair-level accuracy and error bars show 95% confidence intervals. **a**, Cross-scanner retrieval holds the physical section and staining condition fixed while varying the scanner (*n* = 273 slide pairs). **b**, Cross-staining retrieval holds the scanner fixed while comparing co-registered images of serial sections prepared under different staining conditions (*n* = 546 slide pairs). Relative to no normalisation, Reinhard increases Phikon-v2 cross-scanner accuracy but decreases accuracy in the other three encoder–setting combinations; Macenko decreases accuracy in all four combinations.

### 3.2 Exploratory UMAP Shows Partial Scanner-Associated Grouping

To examine whether acquisition conditions were visually associated with the learned representations, we projected the unnormalised Midnight-12k slide-level mean embeddings for all 91 WSIs into two dimensions using UMAP [8] (Fig. 3). This exploratory analysis was restricted to one encoder and preprocessing condition. In the scanner-coloured projection (Fig. 3, panel **a**), AT2, GT450, and P occupied relatively distinct regions, whereas the remaining scanner groups showed substantial overlap. This pattern is consistent with partial scanner-associated grouping in the projection, but does not quantify the scanner variation in the original embeddings space. When the same projection was coloured by staining condition (Fig. 3, panel **b**), the groups were generally dispersed and overlapping. Because each staining condition corresponds to a different serial section in PLISM, differences between staining-condition groups cannot be attributed to colour alone. Overall, the UMAP suggests partial acquisition-associated organisation in this single-model projection. Accordingly, the absence of a consistent retrieval benefit from stain normalisation should not be interpreted as evidence that the encoder is invariant to scanner or staining variation.

**Fig. 3.**
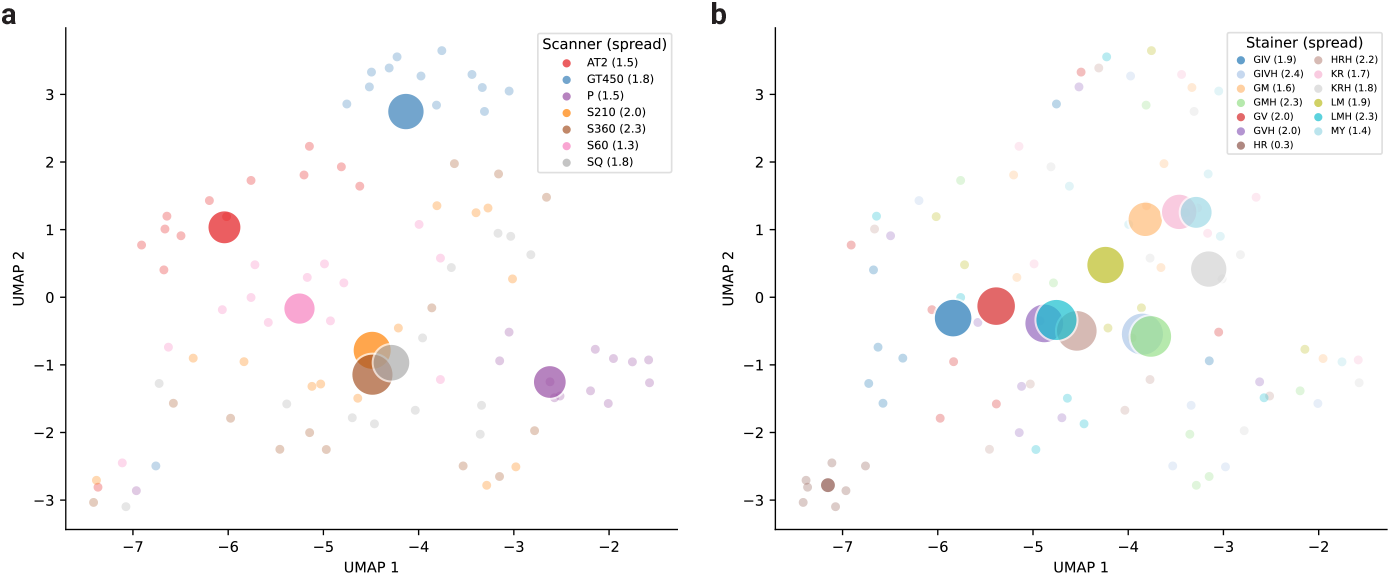
UMAP visualisation of slide-level mean embeddings extracted using Midnight-12k without stain normalisation. Each point represents one of the 91 WSIs, and the same two-dimensional projection is coloured by scanner (panel **a**) and staining condition (panel **b**). Centroid marker size is proportional to within-group spread. Scanner groups show partial separation but substantial overlap, whereas staining-condition groups are generally dispersed and overlapping. Because each staining condition corresponds to a different serial section, panel **b** reflects combined staining- and section-related variation. UMAP was used for qualitative visualisation only.

## 4 Discussion and Conclusion

This study asked whether two commonly used stain normalisation methods improve tile-level retrieval with DINOv2-based pathology encoders under scanner and staining variation. Using the PLISM dataset, we compared Phikon-v2 and Midnight-12k foundation models across three preprocessing conditions (no normalisation, Reinhard, and Macenko) in cross-scanner and cross-staining retrieval settings, and found no consistent benefit from colour normalisation. Reinhard normalisation introduced relatively small, model- and setting-dependent effects, whereas Macenko normalisation generally reduced retrieval performance.

Importantly, the absence of a consistent retrieval benefit should not be interpreted as evidence that the evaluated encoders are colour-invariant or that pretraining conferred robustness to acquisition-related variation. The model- and method-dependent pattern is consistent with the effect of normalisation varying across learned representations. However, the present experimental design cannot isolate the mechanism responsible for these differences. The exploratory UMAP of unnormalised Midnight-12k slide-level embeddings showed partial scanner-associated grouping in two dimensions; however, this single-model projection cannot quantify scanner-associated structure in the original embedding space.

The evaluation was restricted to 400 selected locations from a single tissue microarray block, two DINOv2-based encoders, two normalisation methods, and a single fixed reference tile, limiting generalisability to more heterogeneous datasets, other encoder families, and alternative normalisation or reference-selection strategies. Moreover, cross-staining retrieval compared spatially registered images of serial sections, so staining-related effects could not be separated from section-to-section morphological differences. Finally, the analysis was limited only to tile-level retrieval; these findings should not be assumed to extend to classification, segmentation, or other downstream tasks. Future work should examine multi-institutional datasets, alternative reference-selection strategies, additional encoder architectures and normalisation methods, and test whether the observed pattern extends to slide-level and task-specific evaluations. To conclude, these findings support evaluating stain normalisation empirically for intended encoder, task, and dataset rather than treating it as a default preprocessing step in foundation-model retrieval pipelines.

## Data Availability

The processed PLISM-wsi tiles are publicly available from Owkin at https://huggingface.co/datasets/owkin/plism-dataset-tiles. The analysis code, selected tile identifiers, and pair-level results are available at https://github.com/yaningmeng/stain-normalisation-retrieval.

## Acknowledgments

This work was partly supported by a Google Health unrestricted award to the corresponding author TC at UCL.

